# State-Anxiety Modulates the Effect of Emotion Cues on Visual Temporal Sensitivity in Autism Spectrum Disorder

**DOI:** 10.1101/2021.02.07.430095

**Authors:** Mrinmoy Chakrabarty, Takeshi Atsumi, Ayako Yaguchi, Reiko Fukatsu, Masakazu Ide

**Affiliations:** Dept. of Social Sciences and Humanities, Indraprastha Institute of Information Technology Delhi, New Delhi 110020, INDIA; Dept. of Rehabilitation for Brain Functions, Research Institute of National Rehabilitation Center for Persons with Disabilities, Saitama 359-8555, JAPAN; Dept. of Medical Physiology, Kyorin University, Tokyo 181-8611, JAPAN; Dept. of Psychology, Rikkyo University, Saitama 352-8558, JAPAN

**Author notes:** These authors contributed equally to this research (Mrinmoy Chakrabarty, Takeshi Atsumi, Masakazu Ide).

**Keywords:** emotion, anxiety, visual temporal order judgment, temporal resolution

## Abstract

Atypical processing of stimulus inputs across a range of sensory modalities in autism spectrum disorders (ASD) are widely reported. Sensory processing is known to be influenced by bodily internal states such as physiological arousal and anxiety. Since a sizeable proportion of ASD individuals reportedly have co-morbid anxiety disorders that are linked with dysregulated arousal, we investigated if face-emotion arousal cues, influenced visual sensory sensitivity (indexed by temporal resolution) in an ASD group (n=20) compared to a matched group of typically-developed individuals (TD, n=21). We asked further if emotion-cued changes in visual sensitivity associated with individual differences in state- and trait-anxiety. Participants reported the laterality of the second of two consecutive Gaussian-blob flashes in a visual temporal order judgment task (v-TOJ), demanding higher-level visual processing. The key manipulation was presenting a task-irrelevant face emotion cue briefly at unexpected time points preceding the task-relevant flashes. Disgust vs Neutral emotion signals enhanced the visual temporal resolution in ASD individuals. Furthermore, individual state-anxiety scores correlated with the emotion-cued change of temporal resolution (Disgust vs Neutral) in the ASD group. Both these effects were absent in the TD group. The results show that individual state-anxiety levels significantly modulate the effect of emotions on visual temporal sensitivity in ASD individuals, which was absent in our TD sample. The findings support a nuanced approach to understand the disparate sensory features in ASD individuals, by factoring in the interplay of the individual reactivity to environmental affective information and the severity of anxiety.

## Introduction

Atypical sensory responses across multiple sensory domains are widely reported in autism spectrum disorders (ASD), varying considerably in their manifestations e.g., under-responsiveness, over-responsiveness, fluctuations and unusual interest in sensory aspects of the environment (Baranek et al., 2014; Ben-Sasson et al., 2009). Estimates place the prevalence of sensory abnormalities in ASD between 69 – 93 % (McCormick et al., 2016) and these features are now included as clinical diagnostic criteria of ASD in Diagnostic and Statistical Manual of Mental Disorders, 5th Edition (American Psychiatric Association, 2013). However, the underlying properties of sensory processing in ASD that may explain these unusual behavioral patterns are incompletely understood.

Of the literature pertaining to visual domain in ASD individuals, there are several reports of unusual sensory processing compared to typically developed individuals (TD). Superior processing in visual domain have been reported, such as enhanced identification of simpler luminance-defined gratings but not complex texture-defined ones (Bertone et al., 2005), superior spatial detection of static stimuli (O’Riordan et al., 2001; Pellicano et al., 2005) or pronounced local visual-spatial bias (Jolliffe & Baron-Cohen, 1997; Mottron et al., 1999). In contrast, inferior visual processing have also been demonstrated like impaired object-boundary detection (Vandenbroucke et al., 2008), inferior contrast detection with static and moving stimuli (Sanchez-Marin & Padilla-Medina, 2008), decreased sensitivity to global motion coherence (Spencer et al., 2000) and recognition of biological motion (Annaz et al., 2010; Blake et al., 2003). These studies agree with the notion that lower-level visual processing are generally superior in ASD individuals relative to higher-level, for example human biological motion perception (Dakin & Frith, 2005) and ensemble perception of facial emotions (Chakrabarty & Wada, 2020).

The above studies raise a question whether the heterogeneity of visual sensory characters in individuals with ASD is merely due to individual variations in neural processing of visual information or a resultant of its interaction with additional factors e.g. the internal bodily states such as arousal and anxiety, which to our knowledge has not been clear in these earlier studies. Central and autonomic nervous system responses are widely influenced by arousal through reciprocal noradrenergic projections across prefrontal, sensory and limbic areas, adaptively regulating the responsiveness to sensory inputs by modulating the gain (responsivity) of processing in the task-relevant cortical circuits (Aston-Jones & Cohen, 2005). Indeed, the psychophysical literature with TD individuals shows that arousal evoking emotional content from faces regulate low-level visual perception - precision of orientation discrimination and perceived contrast (Barbot & Carrasco, 2018), orientation sensitivity of low and high spatial frequencies (Bocanegra & Zeelenberg, 2009), contrast sensitivity (Phelps et al., 2006) as well as global motion discrimination (Allen et al., 2016). Emotional faces were also found to facilitate spatial attentional effects on orientation sensitivity (Ferneyhough et al., 2013) and attentional effects on spatial and temporal resolution in gap detection (Bocanegra & Zeelenberg, 2011). The potential role of arousal in visual sensory processing keeps open the question if higher-level visual processing in ASD is also amenable to modulation by environmental emotion cues e.g. facial emotions and if those effects are additionally conditioned by individual background psychopathologies (Yiend, 2010), e.g. anxiety. This is plausible due to the following reasons. Consistent autonomic nervous system differences between ASD and TD individuals, both in sympathetic (Schoen, Miller, Brett-Green, & Nielsen, 2009) and parasympathetic branches of the system (Matsushima et al., 2016; Ming et al., 2005; Schaaf et al., 2003) have related these differences to sensory abnormalities in ASD. This account is supplemented by reports that ~29 – 40 % of ASD population have co-morbid anxiety disorders (Simonoff et al., 2008; van Steensel et al., 2011), which in turn are linked with alterations in arousal (Hoehn-Saric & McLeod, 2000)..

Neural activity of amygdala and its reciprocally connected region - insula have been implicated in anxiety disorders (Gogolla, 2017; Paulus & Stein, 2006; Stein et al., 2007). Moreover, co-occurring anxiety in ASD individuals relates to the activity of amygdala (Herrington et al., 2017), that has projections to sensory processing regions of the brain (LeDoux, 2007). We explored whether temporally unexpected emotion cues below conscious awareness, also processed by the amygdala (Chapman & Anderson, 2012; Phelps & LeDoux, 2005), affected the measures from a task requiring higher-level visual sensory processing in ASD vs. TD individuals. With regards to sensory processing, temporal information is known to play a crucial role (Mauk & Buonomano, 2004) and a recent study has found the severity of sensory manifestations in ASD to correlate with the temporal resolution of stimuli in a tactile TOJ task (Ide et al., 2019). Therefore, we tested whether the temporal resolution of ASD individuals in a visual temporal order judgment task (v-TOJ) demanding higher-level visual processing (Hirsh & Sherrick, 1961; Jaśkowski, 1991) was influenced by facial emotion signals: Disgust (DI) relative to Neutral (NE) [DI>NE]. It is worth mentioning that the study was intended to evaluate the effects of the emotional cues below explicit perceptual judgment and did not require distinguishing the emotions from faces. Facial DI emotion cues were used as DI is entwined with interoceptive processes; elicits physiological arousal regardless of awareness in TD (Allen et al., 2016; Chapman & Anderson, 2012) and also captures visual attention in ASD individuals (Zhao et al., 2016). Yet another reason to investigate the effects of DI in ASD individuals was the fact that core ASD characteristics are linked with social anxiety (Spain et al., 2018) and DI emotional expressions signaling aversion or rejection are rated more negatively than ‘angry’ in socially anxious individuals than non-anxious (Amir et al., 2008), which is of relevance in social communication and relationships (Chapman & Anderson, 2012). Here, it is noteworthy that especially with regards to individuals with ASD, NE may be considered an emotion and not just a null cue (Tottenham et al., 2014). Additionally, we asked whether the influence of emotional signals on the temporal resolution in ASD individuals was modulated by individual state- and trait-anxiety. Effects were compared with a matched TD group. We report marked relative influence of DI to NE emotion signals in enhancing temporal resolution of v-TOJ in ASD individuals and its further modulation by the individual state-anxiety such that temporal resolution worsened with increasing anxiety. Both effects were absent in the TD group.

## Experimental Procedure

### Participants

We recruited 41 Japanese participants - 20 ASD (mean ± standard deviation [s.d.]: age =23.40 ± 5.11 years) and 21 TD (mean ± s.d.: age = 21.09 ± 3.86 years) individuals who were naïve to experimental goals and provided written informed consent to participate in the experiment for a suitable remuneration. These participant numbers were found to be adequate for detecting the effect sizes of interest in the statistical tests with an alpha level of 0.05 and statistical power of 0.80 (see details of a priori sample size calculation in Supplementary Section S1). The ASD and TD groups were matched by gender (Yates’ Chi-square *χ*^*2*^*(1)* = 0.7, *p* = 0.40, *Cramer’s V* = 0.18), age (independent *t*-test: *t*_*(39)*_ =1.63, *p* = 0.11, *Hedge’s g* = 0.51) and full-scale intelligence quotient (independent *t*-test: *t*_*(39)*_ = 1.76, *p* = 0.09, *Hedge’s g* = 0.55). All ASD individuals were recruited from the parent groups of children with developmental disorders and met DSM – IV/IV-TR and/or DSM – 5 criteria of clinical diagnosis by psychiatrists as confirmed from their medical records. The TD individuals were recruited from a pool of university students through social media advertisements and did not report any clinical diagnoses of current or history of psychopathology/ having received any kind of psychological services. The intelligence quotient of the participants assessed by the Japanese version of Wechsler Adult Intelligence-Scale-Third Edition (Japanese WAIS-III Publication Committee, 2006) and other details are in Table 1. Everyone had normal or corrected to normal vision. All procedures were approved by the Ethics Review Board of the institute where the study was carried out.

**Table 1.** Participant information.

| Particulars | ASD | TD |
| --- | --- | --- |
|  | mean (range) | mean (range) |
| Gender ratio (male : female) | 14:6 | 11:10 |
| Age in years | 22.40 (16-33) | 21.09 (16-32) |
| VIQ | 110.7.00 (70-140) | 118.85 (100-136) |
| PIQ | 104.5 (87-122) | 110.33 (80-131) |
| FSIQ | 111.15 (76-134) | 118.48 (93-130) |
| STAI |  |  |
| State-anxiety | 45.20 (31-72) | 39.29 (23-56) |
| Trait-anxiety | 57.00 (39-75) | 42.95 (6-61) |
| VIQ verbal intelligence quotient, PIQ performance IQ, FSIQ full-scale IQ; STAI State Trait Anxiety Inventory |  |  |

### Subjective ratings

All participants completed Japanese versions of the following self-rated measures: State-Trait Anxiety Inventory (STAI) for anxiety levels (Iwata et al, 1998) and the autism spectrum quotient (AQ) for measures of autistic traits (Baron-Cohen et al., 2001; Wakabayashi et al., 2004). Everyone completed the STAI in the rest period between the two counterbalanced blocks of three sessions each (pertaining to one emotion type; see details in ‘*Procedures*’), whereas the AQ was completed just before beginning the experiment.

### Procedures

Participants sat with their heads stabilized using a chin- and head- rest to view stimuli presented on a 27-inch colour monitor (1,920 × 1,080 pixels, 144 Hz; ACER XB27OH) placed at a distance of 60 cm (edges subtending ~ 60 ° × 34°). The experiment was controlled using custom code written in MATLAB 2017a (Mathworks, Natick, Massachusetts, USA) with the Psychophysics Toolbox-3 (Brainard, 1997). Each trial began with a central white fixation cross (1.0° × 1.0°), presented for a random duration between 1,000 and 1,500 ms. The offset of the cross was followed without delay by an image-prime (~ 10° × 10° visual angle), presented at the screen centre for 150 ms. After an intervening blank randomly between 300 and 600 ms, two successive Gaussian-blob patch (sigma value/size = 0.3°; colour = white; luminance = 28.9 cd m^−2^) visual stimuli were flashed at horizontal eccentricities ± 7.5° from the screen centre, for durations of ~ 7 ms (each). The stimulus onset asynchronies (SOAs) between the two stimuli were chosen pseudo-randomly from a set of eight SOAs (−13.8, −20.8, −41.7, −83.3, 13.8, 20.8, 41.7, 83.3 ms; negative intervals indicate that ‘left-sided’ flash was the first of the two) in each trial. One SOA was tested 12 times, each with Face and Scrambled image-prime trials over three sessions in one condition (Neutral and Disgust). The background screen was grey (luminance =14.6 cd m^−2^) throughout. Participants were instructed to maintain fixation at the screen centre throughout the trial, ignore the picture-prime, accurately judge the laterality (left/right) of the ‘last/second’ of two consecutive flashes and indicate their judgment speedily and accurately on a response pad (RB-740, Cedrus Corporation, San Pedro, CA). The trial ended with the keypad response and the next trial began after a blank of 500 ms. There was no emotion recognition step in the trial sequence in order not to explicitly orient the participants to the emotion cues as effect of emotions on visual temporal resolution was studied below ‘awareness’ (explicit perceptual recognition of an emotion). The task structure is illustrated in Figure 1A.

**Figure 1.**
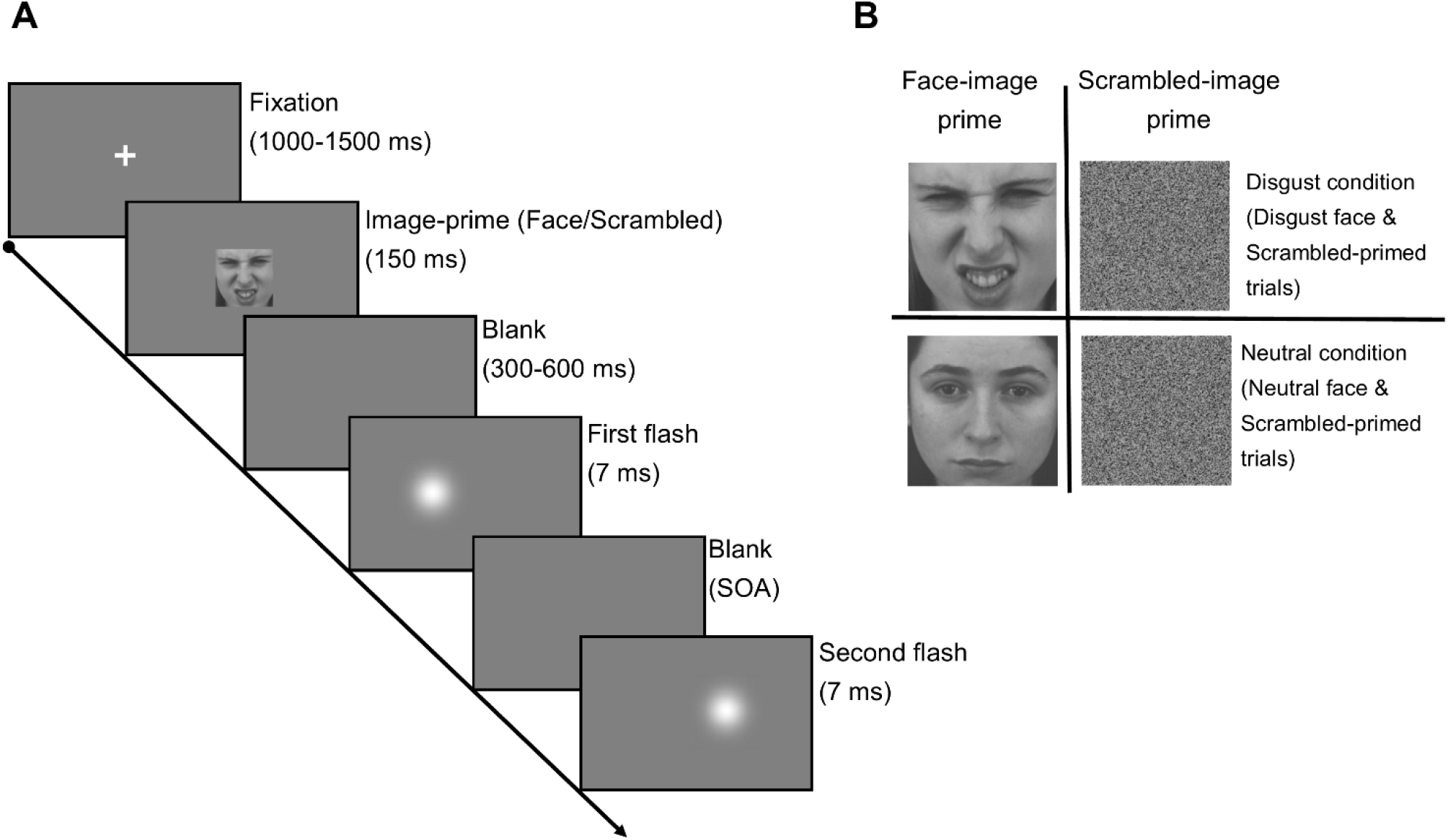
Experimental Procedures. **(A)** The sequence of events in one trial of the visual temporal order judgment task (v-TOJ), pre-cued by an image conveying either emotion (Face) or no information (Scrambled) **(B)** Two different types of experimental conditions based on the emotional valence of the ‘Face’ in the image-prime step of the trials, i.e. ‘Disgust’ condition: Disgust Face- and Scrambled-image trials (32 each × three sessions); ‘Neutral’ condition: Neutral Face- and Scrambled-image trials (32 each × three sessions). KDEF image IDs: top left (Disgust - F05DI) bottom left (Neutral – F15NE). The KDEF images may be included in manuscripts as per the terms of the original authors (https://kdef.se/home/using%20and%20publishing%20kdef%20and%20akdef.html).

Each participant performed six sessions of 64 trials each, with intermittent breaks (total = 384 trials). One session consisted of 32 Face- and Scrambled-trials (control trials) each, which were similar in all respects except that the image-primes in these trials were face and scrambled images (no face/emotion information) respectively. The Face-trials in three of these sessions had Neutral-face images only (hereafter, NE or Neutral-sessions) whereas three others had just Disgust-face images (hereafter, DI or Disgust-sessions) as picture-prime. A block of three sequential sessions of one type (e.g. NE) were followed by a block of three sessions of another type (e.g. DI). The order of the blocks was counterbalanced across participants within a group (ASD/TD). A block of three sequential sessions of one type (e.g. NE) were followed by a block of three sessions of another type (e.g. DI). The order of the blocks was counterbalanced across participants within a group. The main experimental manipulation was the temporally unexpected presentation of the task-irrelevant image-prime (Neutral / Disgust / Scrambled) preceding the actual TOJ task. If emotions influenced visual temporal order processing, then the correct order judgment probability as a function of the SOAs would vary in the trials wherein the image-primes conveyed emotion signals (Neutral / Disgust) compared to the trials with no emotion prime (Scrambled). The experimental condition is illustrated in Figure 1B. Participants performed 10-20 practice trials before commencing the experiment which comprised face image-primes different from experiment trials. During the practice trials an experimenter always confirmed whether the participants were able to distinguish between the different emotions and scrambled cues as well as whether they were able to clearly see the two Gaussian blob flashes on the screen from where the participants were seated.

Face images from academic, non-commercial databases were used: KDEF- Karolinska Directed Emotional Faces database (Flykt et al., 1998) and the NimStim facial expression database (Tottenham et al., 2009). The databases permit using the images for non-commercial academic research. These databases have previously demonstrated effects in ASD (Kanat et al., 2017; Tottenham et al., 2014) and TD individuals (Allen et al., 2016; Greenberg et al., 2017). Two sets of ninety-six images separately expressing Neutral and Disgust emotion (equal number of females and males for each emotion), were selected meaning the overall set comprised 192 images (KDEF: Caucasian ethnicity; NimStim: African-American, Asian-American, Caucasian ethnicities). Thus, different images were presented in all Face-trials (32 images/session × 6 sessions = 192 images). All images were converted to grayscale eliminating the hue and saturation information followed by adjustment of the brightness to the same level (mean = 128, in a range of 0 – 255). Equiluminant scrambled images were separately created from the grayscale images by randomly permuting the coordinates of the pixels so that there was no recognizable face information.

### Data analysis

Keypad response data were analyzed by custom codes. The following explains the analysis steps for a single participant and exemplified in Figure. 2. The response data were sorted by the SOAs to calculate the order judgment probability that the blob on the right side of the fixation cross was flashed later (or the left side of the fixation cross was flashed first). The judgment probabilities were fitted with a cumulative density function of Gaussian distribution (Equation 1) to estimate the temporal resolution (TR) as described earlier (Yamamoto & Kitazawa, 2001). The TR corresponds to the just noticeable difference (JND), which is defined as half of the difference between two SOAs that yielded the judgment that “blob on the right side of the fixation cross was flashed later” in 75% and 25% of the trials (Yamamoto & Kitazawa, 2015).

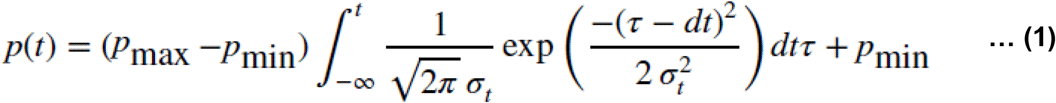

where,

***t, d***_*t*_, ***σ***_*t*_, ***p***_*max*_, ***p***_*min*_ represent the stimulus onset asynchronies, magnitude of horizontal transition, temporal resolution (TR), upper and lower asymptotes of the judgment probability respectively. The TR corresponded to the stimulation interval that yielded 84% correct responses relative to the asymptote. The Optimization Toolbox of MATLAB 2017a (MathWorks, Natick, Massachusetts, USA) was used to minimize the Pearson’s Chi square statistic that reflects the discrepancy between the sampled order-judgment probability and the prediction using the four-parameter model. The model fit was assessed by Pearson’s *r*^2^. As reported in a TOJ study earlier (Spence et al., 2001), participants were included in the final analyses only if the model fit to their data fit attained Pearson’s *r*^2^ ≥ 0.4 and above in all conditions, i.e. in the Disgust condition (three individual model fits to each of the three sessions of Disgust Face-image and Scrambled-image trials) and in the Neutral condition (three individual model fits to each of the three sessions of Neutral Face-image and Scrambled-image trials). By this criterion, 14 ASD and 1 TD participants with Pearson’s *r*^2^ < 0.4 were excluded from the final analyses. The numbers reported under section ‘*Participants*’ reflect the actual number of participants after this exclusion. The mean of Pearson’s *r*^2^ values from each of the Face and Scrambled trials for all the sessions were ≥ 0.85 (see Supplementary Information Table S2).

**Figure 2.**
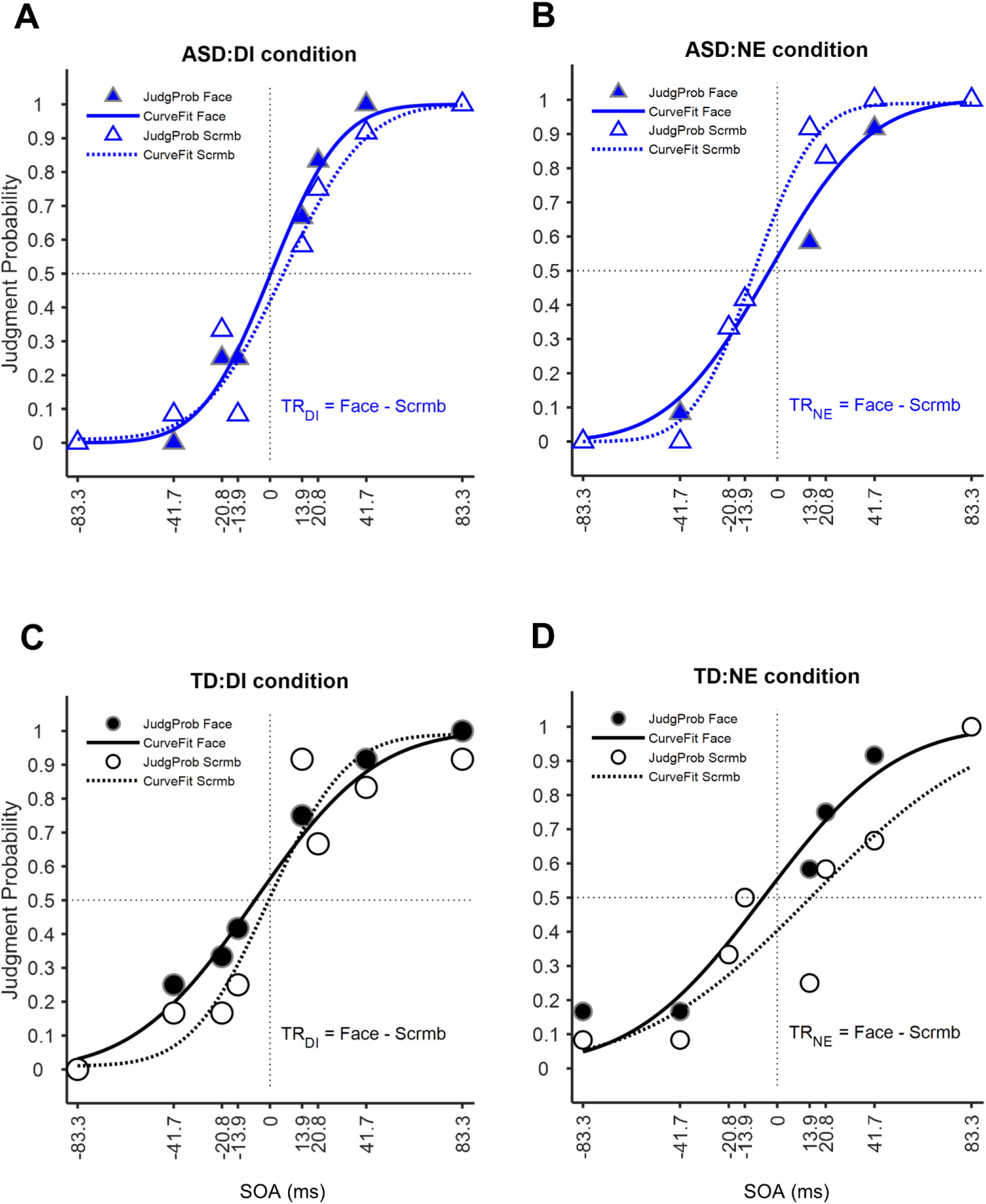
Calculation of visual temporal resolution. **(A-B)** Data from one participant of autism spectrum disorder (ASD). Temporal order judgment probability in the Face-image (filled blue triangles) and Scrambled-image (open blue triangles) trials of Disgust (DI; **A**) and Neutral (NE; **B**) conditions respectively. **(C-D)** Data from one typically-developed participant (TD). Temporal order judgment probability in the Face-image (filled black circles) and Scrambled-image (open black circles) trials of Disgust (DI; **C**) and Neutral (NE; **D**) conditions. The cumulative Gaussian fits (Equation 1) to the order judgment probability (y-axis) as a function of the stimulus onset asynchrony (x-axis) are shown for Face-trials (solid curves) and Scrambled trials (broken curves) in ASD (blue, **A-B**) and TD (black, **C-D**).

Estimates of TR (measure of visual sensitivity) as above were obtained separately for the Face-trials and the Scrambled-trials in each of the three sessions for a condition (DI and NE). Next, the TR of scrambled-trials (control/baseline) was subtracted from that of Face-trials to isolate facial emotion-specific effects on TR after removing the low-level image properties of the pre-cue. This resulted in one TR for each session (NE sessions: TR_NE_; DI sessions: TR_DI_). See Figure 2 for an illustration. Mean of TRs across three NE sessions yielded TR for NE condition. A similar step across three DI sessions yielded TR for DI condition. Finally, TR_change_ (measure of change in visual sensitivity) due to DI emotion signals relative to NE (DI>NE) was calculated as mean (TR_DI_) minus mean (TR_NE_) for each participant, and later used for all statistical analyses. Such subtraction based metrics to isolate target (e.g. face) specific effects and contrasting a dependent measure in two experimental conditions have been reported earlier (Barbot & Carrasco, 2018; Coggan et al., 2017; Kuefner et al., 2010; Rousselet et al., 2008). A significant correlation between variables, i.e. minuend (e.g.TR_DI_) and subtrahend (e.g.TR_NE_) of the subtraction based metric (e.g.TR_change_) is a concern of the utility of such subtraction based metrics. Thus, we separately confirmed that there were no such correlations in our data (see *r* and *p* values in Supplementary Table S3).

Within- and between-group differences were tested by permutation based one-sample and independent-sample *t*-tests respectively which are free from assumptions of underlying distributions of data. Association of state- and trait-anxiety scores with TR_change_ was tested by permutation-based Pearson’s correlation coefficients. For all analyses 20,000 permutations were used to estimate the distribution of the null hypothesis and implemented as described (Groppe et al., 2011) with the Statistics and Machine Learning Toolbox (version 10.0) of MATLAB 2015a. The difference between two independent correlation coefficients was tested using Fisher’s *r*-to-*z* transformation as explained here (Preacher, 2002). For all purposes, *p* < 0.05 was considered statistically significant.

## Results

### Effects of emotion cues on visual temporal resolution

Initial tests done to confirm whether the TR in Scrambled-image trials across DI and NE conditions due to low-level properties of the image-cues varied in the two participant groups revealed no significant difference either between the two conditions within each group (ASD group: DI condition, mean ± sem = 30.19 ± 4.89, NE condition, 24.39 ± 2.87; *t*_*(19)*_ *=* 1.46, two-tailed *p =* 0.16, *Cohen’s d* = 0.56 and TD group: DI condition, 20.10 ± 2.05, NE condition, 24.69 ± 3.02; *t*_*(20)*_ *=* −1.67, two-tailed *p =* 0.11, *Cohen’s d* = 0.39) or within the same condition between the two groups (DI condition: ASD vs. TD group, *t*_*(39)*_ *=* 1.94, two-tailed *p =* 0.06, *Hedge’s g* = 0.60 and NE condition: ASD vs. TD group, *t*_*(39)*_ *=* −0.07, two-tailed *p =* 0.94. *Hedge’s g* = 0.02). Individual TR values of both Face- and Scrambled-image trials before calculating TR_change_ are shown in Figure 3A.

**Figure 3.**
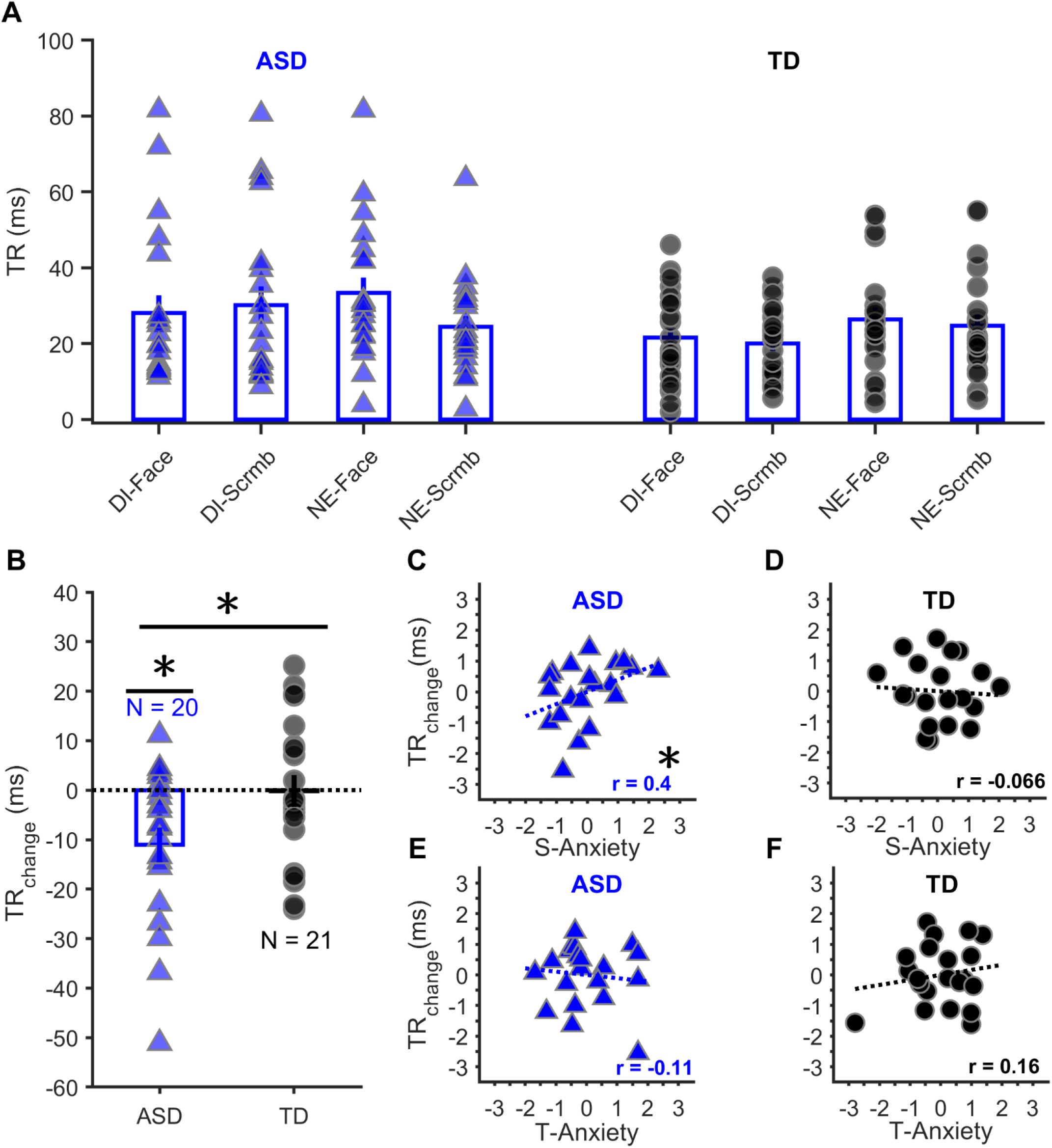
Effects of emotion cues and anxiety on visual temporal resolution. **(A)** Temporal resolution (TR) averaged across trials of individual participants and across participants for each session in ASD (solid blue triangles with bars) and TD (solid black circles with bars) respectively. The sessions were pre-cued by DI-Face (Disgust-face), DI-Scrmb (Disgust-scrambled), NE-Face (Neutral-face), NE-Scrmb (Neutral-scrambled) images respectively. **(B)** Temporal resolution change (TR_change_) of individual participants with group-averaged values in ASD (solid blue triangles with bars) and TD (solid black circles with bars) respectively. Note the trend of overall enhancement of temporal resolution with Disgust relative to Neutral in the ASD group reflected as decrements of TR_change_. **(C-D)** Correlations between the S-Anxiety (State-Anxiety) scores and TR_change_ in ASD (blue, filled triangles and broken line; **C**) and TD (black, filled circles and broken line; **D**). Note a significant correlation in ASD but not TD. (**E-F**) Correlations between the T-Anxiety (Trait-Anxiety) scores and TR_change_ in ASD (blue, filled triangles and broken line; **E**) and TD (black, filled circles and broken line; **F**). Error bars in **(A) and (B)** represent ± 1 SEM; **\****p*<0.05.

The effects of the emotion signals on visual temporal sensitivity within- or between-groups were assessed by TR_change_. Negative TR_change_ value indicates enhancement (resolving finer time intervals between two visual stimuli), whereas a positive value deterioration, as a result of Disgust relative to Neutral emotion-cues (DI>NE). As exemplified in Figure 3B, tests of within-group difference of means from the reference value zero (= no effect of emotion signals) revealed a significant negative TR_change_ in the ASD (mean ± sem = −11.07 ± 3.51; *t*_*(19)*_ = −3.15, two-tailed *p =* 0.003 *, Cohen’s d = 0.70*) but not the TD group (−0.18 ± 3.23; *t*_*(20)*_ *=* −0.06, two-tailed *p =* 0.95*, d = 0.01*), suggesting TR improved with emotion signals only in ASD individuals. Additionally, the size of TR-enhancement in the ASD group was significantly better than TD (*t*_*(39)*_ *=* −2.29, two-tailed *p =* 0.03*, Hedge’s g = 0.72*).

The analyses reported above as separate *t*-tests were further confirmed by a 2 x 2 MIXED ANOVA with factors of Condition (levels: Disgust, Neutral; within-subjects factor) x Group (levels: ASD, TD; between-subjects factor). The conclusions drawn from the results of ANOVA (Supplementary Section S4) agree with the above analyses.

### Modulation of emotion-induced effects on visual temporal resolution by anxiety and autistic traits

The presence of anxiety in the participants was assessed by the STAI instrument. Earlier studies had found scores at or exceeding the threshold of 39-40 on the STAI-state scale corresponded with clinically significant symptoms of a state of anxiety (Addolorato et al., 1999; Forsberg & Björvell, 1993; Knight et al., 1983). Since a cut-off score of 42 on this scale was earlier used to define anxiety disorders in the Japanese population (Kitazawa et al., 2018), which also constituted our samples, we calculated the percentage of individuals in the ASD and TD groups respectively, who had STAI-state scores ≥ 42. By this criterion, we found 60% of ASD and 42.9 % of TD individuals to have anxiety disorders. The state-anxiety scores of the ASD group (mean ± sem = 45.20 ± 2.60) were higher than TD (39.29 ± 1.79) which missed significance marginally (*t*_*(39)*_ *=* 1.89*, p =* 0.07, *Hedge’s g =* 0.59).

First, it was tested if individual state-anxiety intensities modulated the observed effect of emotion signals on TR_change_..The individual state-anxiety ratings of ASD and TD groups were correlated with their respective TR_change_ values. In this regard, a previous study showed that the difference of low-level visual working memory accuracies (Fear trials minus Neutral trials) correlated positively and significantly with state-anxiety in TD individuals (Berggren et al., 2017).Specifically, they found that greater state-anxiety impaired visual processing following Neutral but not Fear cues. Based on this result, we assumed a similar positive association of the state- anxiety scores with our TR_change_ metric (TR_Disgust_ trials minus TR_Neutral_ trials).Since, there were reports that ASD individuals have a higher rate of anxiety than TD (Vasa & Mazurek, 2015), greater proportion of ASD participants were expected to have higher levels of anxiety than TD. Thus, we further assumed that the strength of positive association between state-anxiety and TR change within our ASD group would be higher than our TD group. This correlation revealed significance in the ASD group (blue regression line; *r* = 0.40, one-tailed *p* = 0.036; Figure. 3C), such that the emotion-signalled TR_change_ worsened (due to impairment of higher-level visual processing with NE, reflected by less negative TR_change_ values) with increasing state-anxiety in ASD individuals. However, such association was insignificant in the TD group (black regression line; *r* = −0.07, one-tailed *p* = 0.61; Figure. 3D). The strength of association was greater in the ASD group than TD such that the difference between the two independent correlation coefficients attained a medium effect size (Cohen, 1988), but could not reach significance (details in *Data analysis*; *z* score = 1.46, one-tailed *p* = 0.07, Cohen’s *q* = 0.49).

On the other hand the trait-anxiety intensities as indicated by the respective scores of ASD individuals (mean ± sem = 57 ± 2.39) were much higher than TD individuals (42.95 ± 2.88) which was significant (*t*_*(39)*_ *=* 3.73, two-tailed *p =* 0.00061, *Hedge’s g =* 1.17). Next, associations between trait-anxiety of neither the ASD nor the TD group and their respective TR change values were significant (ASD: blue broken line, *r =* −0.11, two-tailed *p =* 0.63, Figure 3E; TD: *r =* 0.16, two-tailed *p =* 0.48; Figure 3F).

Finally, the measure of autistic traits (AQ score) in ASD individuals (mean ± sem = 33.80 ± 1.69) significantly exceeded that of TD individuals (21.05 ± 1.46; *t*_*(39)*_ *=* 5.71, two-tailed *p =* 1.3 × 10^−6^, *Hedge’s g =* 1.78), but scores of neither group associated with their respective TR_change_ values (*r* and *p* values in Supplementary Table. S5).

## Discussion

We examined how facial emotion cues of Disgust (DI) relative to Neutral (NE), influenced the visual temporal resolution (TR; index of visual temporal sensitivity) in ASD and TD. Consequently, we found that DI vs NE emotion cues markedly enhanced the temporal sensitivity in ASD compared to TD individuals. Furthermore, individual state-anxiety scores significantly modulated the temporal sensitivity in ASD individuals. In contrast, there was neither an apparent emotion-induced effect on temporal sensitivity nor any modulatory trend of the same by state-anxiety in TD individuals. Collectively, the findings reveal that the effects task-irrelevant visual affective signals have on high-level visual temporal processing associate with individual state-anxiety levels in individuals with ASD which is not evident in the TD individuals of our sample.

We found no difference between the temporal resolutions of ASD and TD groups with emotion-null, pre-cues (Scrambled-image trials) that differed from an earlier visual simultaneity judgment (SJ) study (Falter et al., 2012) showing enhanced temporal resolution in ASD vs TD group. Apart from their un-cued task design and their estimates of perceptual threshold against our just noticeable difference (JND) as index of temporal resolution, the disparity between these findings could also be due to the variant higher-order decision making mechanisms between SJ and TOJ. Jaskowski (1991) hypothesized that there are two-stages of temporal processing for sensory inputs: first-stage generates judgements irrespective of the synchronicity of sensory stimuli; second-stage orders those stimuli. Successful performance in TOJ demands processing in both stages, whereas SJ can be solved just by the first stage (Hirsh & Sherrick, 1961; Jaśkowski, 1991).

Results of ASD individuals’ sensitivity in other sensory domains have also been inconsistent. For instance, in unilateral tactile-TOJ, the TR of ASD individuals with index and middle fingers was better than TD individuals but not bilaterally (Tommerdahl et al., 2008). Another study on bilateral tactile-TOJ found TR of ASD children to be better than TD which was statistically insignificant owing to the large inter-individual differences of the ASD group (Wada et al., 2014). More recently, we found no such difference between the two groups in adults (Ide et al., 2019). In all these studies including the former (Falter et al., 2012), the internal state of the participants was experimentally unperturbed. Importantly however, recent studies have reported basal (not task-evoked), autonomic dysfunction in ASD individuals (Anderson et al., 2013; Matsushima et al., 2016; Ming et al., 2005), demonstrating weaker correlation between resting-state autonomic measure (electro-dermal response) and neural activities of anterior cingulate and insular cortices as well as weaker connectivity of the default mode network, compared to TD (Eilam-Stock et al., 2014). Here, the task-evoked autonomic dysregulation with stimuli of social/emotional relevance in ASD individuals is noteworthy (Hirstein et al., 2001; Kylliäinen & Hietanen, 2006; Van Hecke et al., 2009). Additionally, a considerable proportion of ASD individuals have comorbid anxiety disorders (Simonoff et al., 2008; van Steensel et al., 2011), in turn associated with altered arousal (Hoehn-Saric & McLeod, 2000) and dysfunctional emotion regulation (Amstadter, 2008). In this backdrop and given the regulatory role of autonomic arousal within a number of neural circuits (Aston-Jones & Cohen, 2005) that integrate perceptual and interoceptive information (Critchley & Harrison, 2013; Singer et al., 2009), we investigated the effect induced affective state has on a higher-order cognitive function (temporal sensitivity of visual sensory inputs) in ASD individuals. We addressed this by directly manipulating participants’ arousal (by external arousal eliciting cues) and measured the effects of this manipulation on TR (e.g. JND).

Our results demonstrated that DI>NE emotional cues significantly enhanced the TR of ASD individuals, thereby clarifying the difference from TD individuals. Importantly, this was apparent after discounting the effects due to low-level image information, ascribing the overall differential effects within and between the two groups to the high-level, emotion information from the Face-image trials. These results support the explanation that the relative extents of cognitive/attentional benefits Disgust confer over Neutral emotion signals influence the temporal sensitivity in v-TOJ. A key brain structure for processing social-emotional information, is the amygdala (Fitzgerald et al., 2006), which shows varying levels of activity based on context and individual differences (Adolphs, 2010). Accordingly, the emotion information (Disgust relative to Neutral) in our task could have been processed differently by the amygdala in the ASD vs TD group. This concurs with reports that amygdala functional activity (Tottenham et al., 2014) and arousal responses (Mathersul et al., 2013), especially to NE stimuli are clearly deviant in ASD individuals. Recently, anxiety in the context of ASD individuals was shown to be related to increased right amygdala activity, when peripherally presented with social visual information and it has been suggested that anxious ASD individuals may perceive neutral facial faces as more anxiogenic by interpreting neutral information as more negative (Herrington et al., 2017). Considering major inter-connections of amygdala with brain areas serving sensory, polymodal association, attention and arousal functions (LeDoux, 2007), it is plausible that the relative patterns of amygdala activity driven by the socially-salient information from faces (Disgust vs Neutral) varied between ASD and TD individuals that differently modulated the distributed information processing in the above network, thereby resulting in our inter-group TR_change_ differences. The above speculation of the influence of amygdala on the inter-group TR_change_ differences manifest in our results, however, awaits further confirmation by direct investigation of brain function.

Our finding - emotion induced TR enhancement in v-TOJ contrasts with a recent report of no apparent emotional benefits to temporal bisection task in ASD (Jones et al., 2017). This might be due to varying processing demands underlying the two tasks in that TOJ tests existence of a ‘perceptual moment’ (Matthews & Meck, 2016), whereas temporal bisection tests subjective perception of the extent/interval of a stimulus/event (Jones et al., 2017). Conversely, the absence of any clear emotion induced effects in our TD sample begs a further question about how the reported benefits of emotions on low-level visual tasks (Barbot & Carrasco, 2018; Bertone et al., 2005; Jolliffe & Baron-Cohen, 1997; O’Riordan et al., 2001; Sanchez-Marin & Padilla-Medina, 2008; Vandenbroucke et al., 2008) extend to those demanding more involved, higher-level visual processing.

Additionally, our results indicated that the extent of individual reactivity to the sudden onset of emotion cues, conditioned by the respective state-anxiety level determined the temporal sensitivity of v-TOJ in ASD individuals. Anxiety is the apprehensive internal/external anticipation of future threat based on no clear object and state-anxiety refers to such a situational, transient emotional and physiological state (Wiedemann, 2015), which is assessed by the state-anxiety subscale of STAI (Spielberger, C.D., Gorsuch, R.L., Lushene, R., Vagg, P.R., Jacobs, 1983). Conforming with suggestions that affective stimuli may lead to anxiety response (Hubert & de Jong-Meyer, 1990; Turner et al., 1991), our results illustrated that emotion-cued changes of temporal sensitivity were predicted by the individual state-anxiety level in ASD individuals, such that the increase of the TR_change_ metric (worsening TR) scaled with the subjective state-anxiety (less negative y-axis values, Fig. 3C). Specifically, the size of NE-induced deterioration exceeded the relative DI-induced enhancement of TR, worsening the overall TR_change_ with increasing state-anxiety.

We found no association between state- or trait-anxiety and v-TOJ task performance in TD individuals, contrary to a recent study that reported emotion × trait-anxiety interaction influences low-level visual task performance (Barbot & Carrasco, 2018). Further, the results from our TD sample showed null effects of emotions on visual processing as against earlier reports of significant effects of emotions on TD visual processing. The reasons for these discrepancies could be the following. First, Barbot & Carrasco (2018) used a delay of 120 ms (measured from the emotional pre-cue onset) to elicit the maximum effect of exogenous attention on the visual task. The earlier reported studies have all used shorter delays of 100-150 ms between the emotion cues and the task-relevant stimuli (Bocanegra and Zeelenberg 2009; Phelps et al 2006; Ferneyhough 2013; Bocanegra and Zeelenberg 2011). By contrast, a similar delay in our task design was much greater (≥ 450 ms), which likely suppressed the effect of emotions on TR_change_ in the TD group. This reason gains support from their results that showed the vanishing effect of emotions when the delay was increased to 500 ms (Barbot & Carrasco, 2018). Second, while all the earlier cited studies have reported the effects of emotion cues on low-level visual tasks in TD individuals (Barbot & Carrasco 2018; Bocanegra and Zeelenberg 2009; Phelps et al 2006; Ferneyhough 2013; Bocanegra and Zeelenberg 2011), we investigated the effects of emotion cues on a high-level visual task. Thus, a mismatch of the levels of visual processing as engaged by the task design could have led to the disparate results in respective TD samples. Together, these accounts encourage a viewpoint that a better understanding of the heterogeneity of task-structure (low-level/high-level visual processing, interval between emotion cue and task stimulus onsets) and study samples (ASD/TD), along with the individual participant’s internal states may be necessary to explain the emotional influences on visual processing.

Finally, we identify some limitations of our study. *First*, we did not objectively measure the autonomic arousal that the emotion cues may have evoked, which may be addressed in future to directly relate the emotion evoked TR changes to the participants’ autonomic state. *Second*, the role of visuospatial attention on the observed effect is not evident in the present study which may be explored, for example by incorporating an eye tracking measure. *Third*, our sample size was relatively modest and a larger study to replicate our findings is warranted.

These limitations notwithstanding, our results clearly demonstrate that temporally unexpected, social affective information influence higher-order visual processing in ASD individuals, which is conditioned by individual state-anxiety levels. The results support a nuanced approach to the disparate sensory features in ASD individuals, by factoring in the interplay of the individual reactivity to affective information and severity of anxiety.

## Acknowledgments

The authors are grateful to all participants for taking part in the experiment. We thank Y. Wang (Clinical Psychologist) for the detailed IQ assessments and Dr. M. Wada (Section Chief) for his overall support.

## Authors’ contributions/credit

**Mrinmoy Chakrabarty:** Conceptualization, Methodology, Investigation, Software, Visualisation, Writing-Original draft preparation. **Takeshi Atsumi:** Conceptualization, Methodology, Investigation, Writing-Reviewing and Editing. **Reiko Fukatsu:** Conceptualization, Supervision, Writing-Reviewing and Editing. **Masakazu Ide:** Conceptualization, Methodology, Investigation, Funding Acquisition, Project Administration, Writing-Reviewing and Editing.

All authors approved the final version of manuscript.

## Competing interests

The authors declare no competing interests.

## Data accessibility

All data that led to the reported results can be accessed from the corresponding author upon a reasonable request.

## Abbreviations

ASD: Autism Spectrum Disorders individuals
TD: Typically Developed individuals
TOJ: Temporal Order Judgment
v-TOJ: visual Temporal Order Judgment
TR: Temporal Resolution
STAI: State-Trait Anxiety Inventory

## Funding

The research was funded by a Grant-in-Aid from Japan Society for the Promotion of Science (grant numbers JP18H03140, JP18H03663, JP18K18705), Ministry of Education, Culture, Sports, Science and Technology-JAPAN (grant numbers JP17H05966) awarded to Masakazu Ide.

## Supplementary Information

### Section S1 A priori sample size calculation

A. *A priori sample size calculation for the differences in TR*_*change*_ *between two independent groups – ASD vs.TD* An earlier study by (Falter et al., 2012) had found a difference of visual temporal resolution between ASD (n = 16) and TD (n = 16) in a task related to TOJ - simultanaeity judgment with no emotional pre-cues. The authors performed a Mann Whitney U test to test the difference between the groups and reported an effect size of 0.4 (*r* = z value of Mann Whitney U / square root of number of individuals) or an equivalent Cohen’s *d* = 0.9 or ~ 1 (Cohen, 1988; Lenhard & Lenhard, 2016), suggesting a large effect. Given this effect size and our emotion pre-cues in the task design, we anticipated a large effect (Cohen’s *d* = 1) between our ASD and TD groups and calculated the sample sizes for a difference between two independent groups by a 2-tailed independent *t*-test with the following particulars…
  A. H1 (alternative hypothesis) = There is a difference between the means of ASD and TD groups.
  B. alpha level (probability of type-I error) = 0.05
  C. power (% chance of correctly rejecting the null hypothesis, H0) = 0.80 (80%)
  D. allocation ratio (sample size of group 2 [TD] / sample size of group 1 [ASD]) = 1.1 (22 TD/20 ASD; the participant numbers were based on the readily available participant pool at the time of initiating the experiment and our expectation of ~10% participant exclusion due to not meeting the objective criterion of Pearson’s *r* of model fit (see Data analysis), based on earlier experience with the temporal order judgement task in our lab.
  E. H0 (null hypothesis) = There is no difference between the means of ASD and TD groups.
  Using the above values in the program G*power, version 3.1.9.7 (Faul et al., 2009), we got sample sizes of at least 16 individuals in the ASD group and 18 individuals in the TD group respectively, to correctly reject the null hypotheses in the 2-tailed independent *t*-test with 80% chance.
B. *A priori sample size calculation for the correlation between State-anxiety score and TR*_*change*_ *in the two independent groups – ASD and TD* An earlier study conducted by (Berggren et al., 2017) with TD participants (N = 42), found that the difference of low-level visual working memory accuracies (Fear trials minus Neutral trials) correlated positively and significantly with state-anxiety (*r* = 0.37 or *r* ~ 0.4) with a 2-tailed test. Specifically, they reported: “state anxiety was associated with impaired accuracy following a neutral cue, with this effect not present following threat.” Based on the above, we assumed a similar positive association (1-tailed hypothesis) of the state-anxiety scores with our TR_change_ metric (TR_Disgust_ trials minus TR_Neutral_ trials). The strength of correlation mentioned above (*r* = 0.4) indicated a conventional effect size between medium to strong (Cohen, 1988). In the Berggren et. al. 2017 study, the sample was from the Western population and had 4.3% (6/42) of the individuals with state-anxiety score ≥ 39-40, which is an accepted threshold that corresponds with clinically relevant high anxiety (Addolorato et al., 1999; Forsberg & Björvell, 1993; Knight et al., 1983).However, our samples were to be drawn from the Japanese population who were reported to have a higher level of anxiety (Iwata et al., 1998) than the Western population. Thus, we assumed a stronger correlation than that reported by Berggren et. al. 2017, i.e., *r* > 0.4 in both our TD and ASD groups. Besides, there were reports that ASD individuals have a higher rate of anxiety than TD (Vasa & Mazurek, 2015) which suggested that a greater proportion of ASD participants were expected to have higher levels of anxiety than TD. Therefore, we further assumed that the correlation within our ASD group would be slightly higher than our TD group. Taken together, in the a-priori sample size calculation for the correlation analysis between state-anxiety and TR_change_, we used the following…
  A. H1 correlation coefficient (alternative hypotheses) - ASD = 0.65; TD = 0.60
  B. alpha level (probability of type-I error) = 0.05 (for both ASD and TD)
  C. power (% chance of correctly rejecting the null hypothesis, H0) = 0.80 (80%; for both ASD and TD)
  D. correlation coefficient assuming null hypothesis (H0) = 0 (for both ASD and TD)
  Using these values in the program G*power version 3.1.9.7, (Faul et al., 2009), we got sample sizes of at least 13 individuals in the ASD group and 15 individuals in the TD group respectively, to correctly reject the null hypotheses in the simple linear regression analyses with 80% chance.

**Table S2.** Pearson’s *r*^*2*^ values of the model fit (Equation 1).

| Condition | Face<br>(Session1) | Scrambled<br>(Session1) | Face<br>(Session2) | Scrambled<br>(Session2) | Face<br>(Session3) | Scrambled<br>(Session3) |
| --- | --- | --- | --- | --- | --- | --- |
| <u>ASD (N = 20)</u> |  |  |  |  |  |  |
| Disgust | 0.87 ± 0.13<br>[0.68, 1] | 0.82 ± 0.10<br>[0.4, 1] | 0.90 ± 0.07<br>[0.63, 1] | 0.85 ± 0.09<br>[0.4, 1] | 0.88 ± 0.15<br>[0.4, 1] | 0.89 ± 0.12<br>[0.74, 1] |
| Neutral | 0.85 ± 0.08<br>[0.5, 0.99] | 0.89 ± 0.06<br>[0.69, 0.98] | 0.85 ± 0.07<br>[0.5, 0.99] | 0.87 ± 0.09<br>[0.81, 1] | 0.86 ± 0.09<br>[0.69, 0.99] | 0.89 ± 0.06<br>[0.76, 1] |
| <u>TD (N = 21)</u> |  |  |  |  |  |  |
| Disgust | 0.91 ± 0.03<br>[0.76, 1] | 0.91 ± 0.04<br>[0.51, 1] | 0.90 ± 0.04<br>[0.64, 1] | 0.88 ± 0.06<br>[0.66, 1] | 0.91 ± 0.07<br>[0.43, 1] | 0.95 ± 0.03<br>[0.78, 1] |
| Neutral | 0.88 ± 0.03<br>[0.6, 0.99] | 0.87 ± 0.05<br>[0.5, 1] | 0.89 ± 0.03<br>[0.66, 1] | 0.89 ± 0.02<br>[0.73, 1] | 0.89 ± 0.04<br>[0.5, 0.99] | 0.89 ± 0.05<br>[0.66, 1] |

Each cell shows the session-wise mean ± standard error of mean values across participants (N) in each group (ASD and TD). Values within square brackets in each cell indicate the respective ranges.

**Table S3.** Pearson’s *r* and *p* values* of the correlation between the minuhend (TR_DI_) and subtrahend (TR_NE_) of TR_change_ metric.

| Correlation<br>between | ASD<br>(N=20) | TD<br>(N=21) |
| --- | --- | --- |
| TR<sub>DI</sub> and<br>TR<sub>NE</sub> | $r = 0.22$;<br>$p = 0.36$ | $r = -0.09$;<br>$p = 0.71$ |
| TR<sub>face</sub> and<br>TR<sub>scrambled</sub><br>(DI session) | $r = -0.36$;<br>$p = 0.15$ | $r = 0.08$;<br>$p = 0.73$ |
| TR<sub>face</sub> and<br>TR<sub>scrambled</sub><br>(NE session) | $r = 0.21$;<br>$p = 0.37$ | $r = -0.38$;<br>$p = 0.09$ |

\* Permutation-based Pearson's correlation coefficients (20,000 permutations, two-tailed $p$ values; details in *Data Analysis*).

### Section S4. Effects of emotion cues on visual temporal resolution: results of ANOVA

To ascertain whether the temporal resolution (TR) in Scrambled-image trials across Disgust (DI) and Neutral (NE) conditions due to low-level properties of the image-cues varied in the two participant groups, a 2 x 2 MIXED ANOVA with factors of Condition (levels: DI, NE; within-subjects factor) x Group (levels: ASD, TD; between-subjects factor) was run on the TR. Consequently, no significant interaction between the factors of Condition and Group (*F*_*(1,39)*_ = 3.53, *p* = 0.07, partial *η*^*2*^= 0.08) was found. There was also no significant within-subject main effect of Condition (*F*_*(1,39)*_ = 0.31, *p* = 0.58, partial *η*^*2*^= 0.01) or between-subject main effect of Group (*F*_*(1,39)*_ = 2.04, *p* = 0.16, partial *η*^*2*^= 0.05). These results suggest that there were no significant changes in the TR of the Scrambled image trials across the whole sample.

Next, another 2 x 2 MIXED ANOVA with the same factors of Condition (levels: DI, NE; within-subjects factor) x Group (levels: ASD, TD; between-subjects factor) was run on the TR data from Face-image trials after discounting the effects of low-level image properties from Scrambled-image trials so as to isolate emotion specific TR changes in the participants. The results revealed a significant interaction between the factors of Condition and Group (*F*_*(1,39)*_ = 5.23, *p* = 0.028, partial *η*^*2*^= 0.12). This suggests that changes in TR across the conditions (DI and NE) are not equivalent across the two groups (ASD and TD). The significant interaction when followed up by pairwise comparisons (simple main effects) revealed that there was a significant difference between DI and NE conditions in the ASD group (difference of means = −11.07 ± 3.41, Bonferroni corrected *p* = 0.002) but not in the TD group (difference of means = −0.18 ± 3.27, Bonferroni corrected *p* = 0.96).

The above results agree with the conclusions drawn from Figure 3A and 3B in the main text.

**Table S5.** Pearson’s *r* and *p* values^*^ of the correlations between the total Autism Spectrum Quotient (AQ) of the ASD and TD groups with their respective TR_change_ values.

| Total<br>AQ | ASD<br>(N=20) | TD<br>(N=21) |
| --- | --- | --- |
| Score | $r = -0.12$;<br>$p = 0.67$ | $r = -0.16$;<br>$p = 0.47$ |

\* Permutation-based Pearson's correlation coefficients (20,000 permutations, two-tailed $p$ values; details in *Data Analysis*).

## Notes

### Competing Interest Statement

The authors have declared no competing interest.

